# Epigenetic regulation of protein translation in *KMT2A*-rearranged AML

**DOI:** 10.1101/790287

**Authors:** Alexandra Lenard, Hongbo Michael Xie, Simone S. Riedel, Zuo-Fei Yuan, Nan Zhu, Tobias Neff, Kathrin M. Bernt

## Abstract

Inhibition of the histone methyl-transferase DOT1L (KMT4) has shown encouraging activity in preclinical models of *KMT2A* (*MLL)*-rearranged leukemia. The DOT1L inhibitor pinometostat (EPZ5676) was well tolerated in early phase clinical trials and showed modest clinical activity, including occasional complete responses (CRs) as single agent. These studies support the development of combinatorial therapies for *KMT2A*-rearranged leukemias. Here, we investigated two novel combinations: dual inhibition of the histone methyltransferases DOT1L and EZH2, and the combination of a DOT1L inhibitor with the protein synthesis inhibitor homoharringtonine (HHR).

EZH2 is the catalytic histone methyltransferase in the polycomb repressive complex 2 (PRC2), and inhibition of EZH2 has reported preclinical activity in *KMT2A*-rearranged leukemia. We found that the H3K79 and H3K27 methyl marks are not dependent on each other, and that DOT1L and EZH2 inhibition affect largely distinct gene expression programs. In particular, the KMT2A/DOT1L target HOXA9, which is commonly de-repressed as a consequence of PRC2 loss or inhibition in other contexts, was not re-activated upon dual DOT1L/EZH2 knockout or inhibition. Despite encouraging data in murine KMT2A-MLLT3 transformed cells suggesting synergy between DOT1L and EZH2 inhibition, we found both synergistic and antagonistic effects on a panel of human KMT2A rearranged cell lines. Combinatorial inhibition of DOT1L and EZH2 is thus not a promising strategy. We identified opposing effects on ribosomal gene transcription and protein translation by DOT1L and EZH2 as a mechanism that is partially responsible for observed antagonistic effects. The effects of DOT1L inhibition on ribosomal gene expression prompted us to evaluate the combination of EPZ5676 with a protein translation inhibitor. EPZ5676 was synergistic with the protein translation inhibitor homoharringtonine (HHR), supporting further preclinical/clinical development of this combination.

## INTRODUCTION

Rearrangements of *KMT2A (MLL1)* occur in 10% of AML and ALL, and 70% of infant ALL. With few exceptions, *KMT2A* rearrangements are associated with a poor prognosis, and *KMT2A-* rearranged *(KMT2A-r)* leukemias have been the target of substantial drug development efforts and clinical research without much impact yet on survival [1]. *KMT2A* fusions are strongly transforming [2], likely due to the profound epigenetic changes they induce. Key histone lysine methyltransferases (KMTs) involved in KMT2A-fusion mediated leukemogenesis are the H3K79 KMT4 (hereafter referred to as DOT1L) [3-6] and the H3K27 KMT6 (hereafter referred to as EZH2) [7-11]. Both have been proposed as therapeutic targets, and pharmacologic inhibitors for both are currently in clinical trials (although EZH2 inhibitors are not being studied in AML).

Direct genomic targets of the KMT2A fusion display aberrantly high H3K79 methylation levels compared with other highly expressed loci [3], and knockdown, knockout, or pharmacologic inhibition of DOT1L results in the transcriptional downregulation of fusion target gene expression in *KMT2A* fusion leukemia models, cells lines and patient samples [3-6]. These findings formed the basis for two phase I/II clinical trials with the DOT1L inhibitor pinometostat (EPZ5676), as well as ongoing combination trials with hypomethylating agents and chemotherapy. Pinometostat was well tolerated and induced single agent responses including two CRs, one of them durable for many months. However, a substantial number of patients failed to respond, and in others, resistance rapidly developed [12]. Several mechanisms of resistance have been reported in model systems since the initiation of these trials. These include classic drug efflux pump mechanisms (ABCB1A/MDR-1/PGP) [13], epigenetic resistance through downregulation of SIRT1/SUV39 [14], and independence of H3K79 methylation through unknown mechanisms [13]. Nevertheless, the clinical experience provided proof of concept that DOT1L inhibition had activity in *KMT2A-r* disease, with largely non-overlapping toxicities with other agents. A critical next question is whether synergistic combinations with other agents can be identified that ultimately improve outcomes for a larger number of patients.

In this study, we investigated the interplay between DOT1L and EZH2 in *KMT2A-r* AML and ALL cells. Inhibitors of both, DOT1L and EZH2 have been reported to have single agent efficacy in models of *KMT2A-r* leukemia, with different mechanisms of action [1, 3-11]. This raises the question whether dual inhibition could be additive or synergistic. At the same time, potential interplay on several levels could also result in antagonistic effects of dual DOT1L/EZH2 inhibition: DOT1L inhibition results in downregulation of direct KMT2A targets such as the later HOXA cluster genes. During embryonic and hematopoietic development as well as in several subtypes of leukemia, the HOXA cluster is silenced and acquires H3K27 tri-methylation by the polycomb repressive complex 2 (PRC2) [15]. Loss of PRC2 function has been linked to increased HOXA cluster expression in leukemia [16, 17]. Inhibition of EZH2, the main KMT of the PRC2 complex, might thus interfere with the silencing of direct KMT2A fusion targets in a manner similar to SIRT-1/SUV39 [14]. Furthermore, AF10, a core member of the DOT1L complex and rate limiting co-factor for the di- and tri-methylation of H3K79 [18], was reported to bind unmodified, but not H3K27 methylated H3K27 [19]. Inhibition of H3K27 methylation might thus increase the ability of AF10 to bind to KMT2A fusion targets and increase H3K79 methylation, resulting in greater difficulty to achieve profound inhibition of H3K79 methylation. In this study, we set out to answer the following questions: Does EZH2 inhibition modulate H3K79 methylation? Is EZH2/PRC2 required for the silencing of KMT2A fusion direct target genes upon DOT1L inhibition? Is dual DOT1L/EZH2 inhibition synergistic or antagonistic in *KMT2A*-fusion driven leukemia?

In brief, we found that the combination of DOT1L /EZH2 inhibition induced pleiotropic and context dependent effects, and did not consistently synergize with each other. However, these studies identified previously underappreciated effects of these inhibitors on protein translation and suggest that protein translation inhibition acts synergistic with DOT1L inhibition.

## METHODS

For primer sequences, antibodies and detailed experimental procedures please refer to the supplemental materials.

### Cell lines

Human leukemia cell lines were obtained from ATCC or DSMZ and maintained in culture as detailed in the supplemental materials. Cell lines were re-authenticated every 6 months in culture.

### Drug Assays

Compounds were dissolved in DMSO, and all dilution series were prepared keeping the DMSO exposure euqal accross all conditions and at <0.1% (<1:1000). Human leukemia cell lines were exposed to EPZ4777, EPZ5676 (DOT1L inhibitor), GSK 126 (EZH2 Inhibitor) or homoharringtonine (HHT) at the indicated concentrations, and cells were replated at equal densities in fresh compound containing media every 3-4 days. Cell growth and viability was assessed either by serial replating and trypan blue exclusion, or XTT assay.

### *Dot1l* and *Ezh2* knockout mice, breeding

Animals were maintained at the Animal Research Facility at the University of Colorado and Boston Children’s Hospital. Animal experiments were approved by the Internal Animal Care and Use Committee. *Dot1l* [3] and *Ezh2 ([7])* conditional knockout mice were previously described and were maintained on a C57BL/6 background.

### Generation of transformed murine cells

Ecotropic retroviral vectors containing murine KMT2A-MLLT3-IRES-GFP, Cre-IRES-pTomato (Cre) and MSCV-IRES-pTomato (MIT) were generated by cotransfection of 293 cells. Lin^-^Sca-1^+^c-Kit^+^ (LSC) cells were transduced with KMT2A-MLLT3-GFP. After 2-7 days, GFP^+^ cells were sorted and transduced with Cre or control. 2-3 days after transduction, cells were sorted and plated in colony assays or liquid culture.

### Biochemical Assays (cell growth, colony growth, apoptosis, cell cycle, western blotting)

For colony assays, sorted transduced cells were plated in methylcellulose M3234 containing IL3, IL6 and SCF at 1000 cells per plate in duplicate, and colonies were scored after 7 days of culture. For liquid culture of murine cells, cells were maintained in media with cytokine support, and replated every 2-3 days. *Dot1l* and *Ezh2* deletion was verified by PCR at each replating beyond day 7. Cell growth and viability were followed by serial cell counts using Trypan blue exclusion. Apoptosis and cell cycle analysis were performed using the Annexin-staining and the Click-IT EdU kit. Protein translation was analyzed using the Click-IT OP-Puro kit. Western blotting for histone modifications was performed on purified histones isolated by acid extraction using the indicated antibodies and controls.

### Histone Mass Spectrometry

Histones were isolated, chemically derivatized and analyzed by mass spectrometry as previously described [20].

### qPCR analysis of HOXA9 and CDKN2A

RNA was isolated from sorted murine transformed progenitor cells or compound treated human cell lines using RNeasy mini columns (Qiagen). Please refer to the supplement for primer sequences. Fold-change of is shown compared vehicle treated control (human cell lines) or wild type control (murine knockout studies).

### Chromatin immunoprecipitation (ChIP)

Chromatin immunoprecipitation for H3K79me2 and H3K27me3 in murine KMT2A-MLLT3 leukemias was performed using rabbit polyclonal antibodies from abcam (ab3594 Cambridge, MA) similarly as described [21].

### RNA amplification and RNA-Seq

RNA was isolated from MV4;11 cells exposed to 7 days of EPZ5676 or GSK126 using RNeasy mini columns (Qiagen). RNA for RNA-Seq was submitted to the UC-Denver genomics core for library preparation and sequencing.

### Data analysis and statistical methods

Histone PTMs were analyzed using the EpiProfile 2.0 computational algorithm [22]. Drug interactions were evaluated for synergy or antagonism using CompuSyn (http://www.combosyn.com/) [23, 24]. RNA-Seq raw Fastq files were aligned using STAR [25] against Human GRCh37 reference and quantified by applying Kallisto [version 0.45.0, PMID: 27043002]. Output from Kallisto was then directly imported into DESeq2 [26] in order to detect differentially expressed genes (DEG). DEGs were deemed as genes with False Discovery Rate (FDR) less than 0.05 level.

### Gene expression data

was deposited at the NCBI Gene Expression Omnibus and is accessible under GSE134369 (https://www.ncbi.nlm.nih.gov/geo/query/acc.cgi?acc=GSE134369).

## RESULTS

### No interplay between H3K27 and H3K79 methylation

In order to assess whether inhibition of EZH2 affects H3K79 methylation (or DOT1L inhibition impacted H3K27 methylation), we exposed the *KMT2A* rearranged cell line MV4;11 to small molecule inhibitors of DOT1L or EZH2 (Figure 1A). Western blotting over a wide range of inhibitor doses revealed no changes of H3K79 methylation upon inhibition of EZH2, or H3K27 methylation upon inhibition of DOT1L. Results were confirmed by mass spectrometry (Figure 1B) at a fixed dose in an extended cell line panel. Despite a higher availability of H3K27 unmodified histones that could serve as binding sites for AF10 [19], H3K79 methylation is not increased upon inhibition of EZH2.

**Figure 1:**
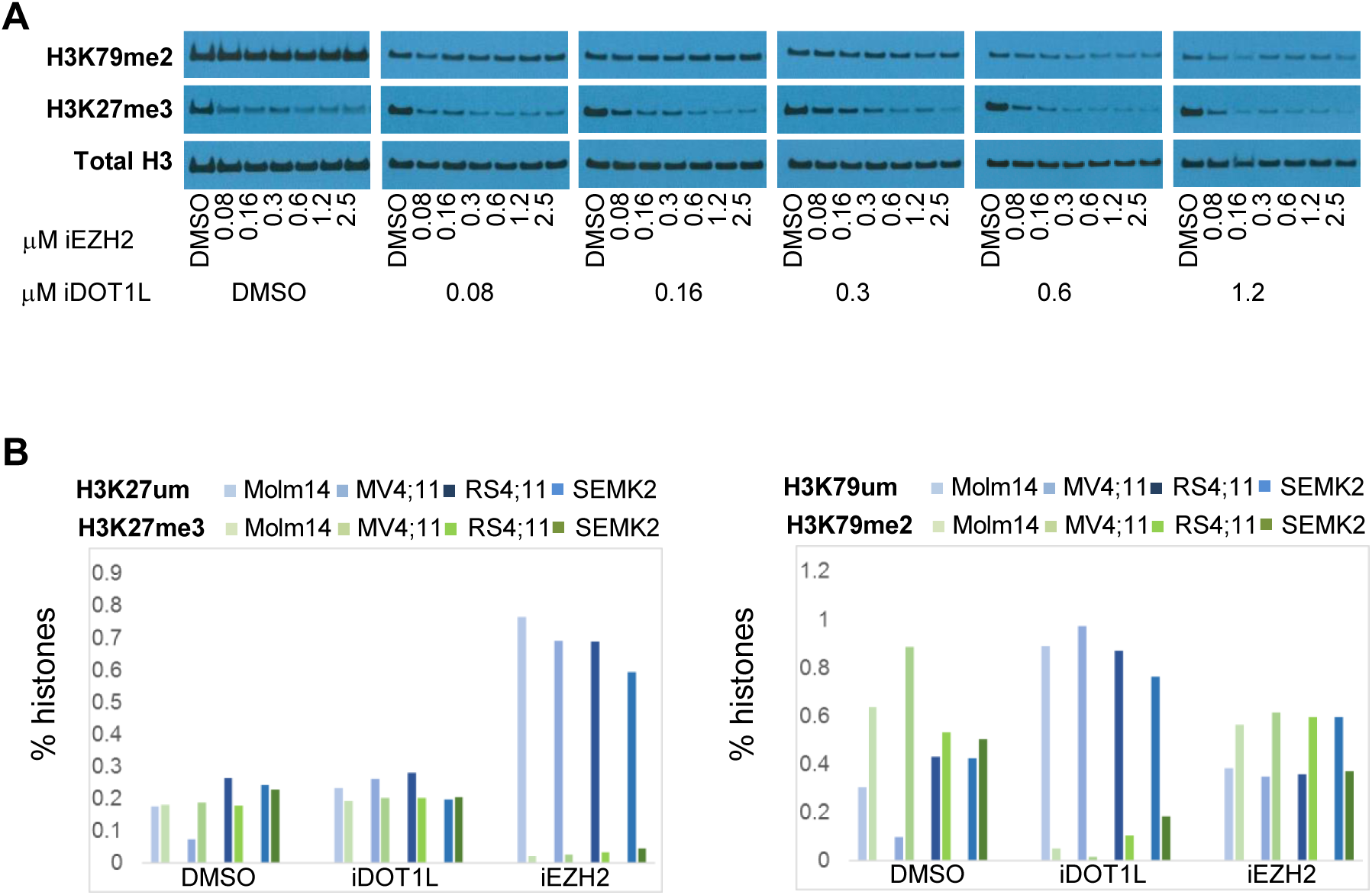
**A:** MV4;11 cells were exposed to the indicated concentrations of EPZ4777 and GSK126 for 4 days, and H3K79me2 / H3K27me3 were detected by Western Blotting. **B:** the indicated cell lines were exposed to 10 μM EPZ5676 (iDOT1L) or 3 μM GSK126 (iEZH2) for 4 days, and H3K79me2 and H3K27me3 were determined by mass-spectrometry.

### Synergy between DOT1L and EZH2 deletion in murine KMT2A-MLLT3 (MLL-AF9) transformed bone marrow cells

We next asked if dual knockout of *Dot1l* and *Ezh2* would act synergistic or antagonistic in a defined genetic mouse model driven by MLL-AF9 (KMT2A-MLLT3). MLL-AF9 was retrovirally introduced into lin-cKit+ Sca-1+ (LSK) cells from mice carrying conditional alleles for *Dot1l* and/or *Ezh2*. Deletion of either Dot1l or Ezh2 alone resulted in smaller and more differentiated colonies (which we previously showed to be devoid of replating capacity, therefore colonies were not replated). Dual inactivation of *Dot1l* and *Ezh2* resulted in a near complete failure to yield any colony formation (Figure 2A), and particularly rapid outgrowth of cells that had failed to undergo complete deletion of the *Ezh2* allele (Figure 2B). Transcriptionally, inactivation of *Ezh2* was accompanied by de-repression of *Cdkn2a*, which was independent of Dot1l and H3K79 methylation. In turn, HoxA9 expression was decreased upon loss of *Dot1l*, which was not rescued by concomitant loss of *Ezh2* (Figure 2C). Consistent with these results, ChIP-qPCR of the later *HoxA* cluster showed no substantial increase in H3K27 trimethylation 5 days after Dot1l deletion, while HoxA9 expression was already profoundly decreased. We therefore conclude that PRC2 is not the primary mechanism by which the HoxA cluster is silenced upon loss or inhibition of DOT1L.

**Figure 2:**
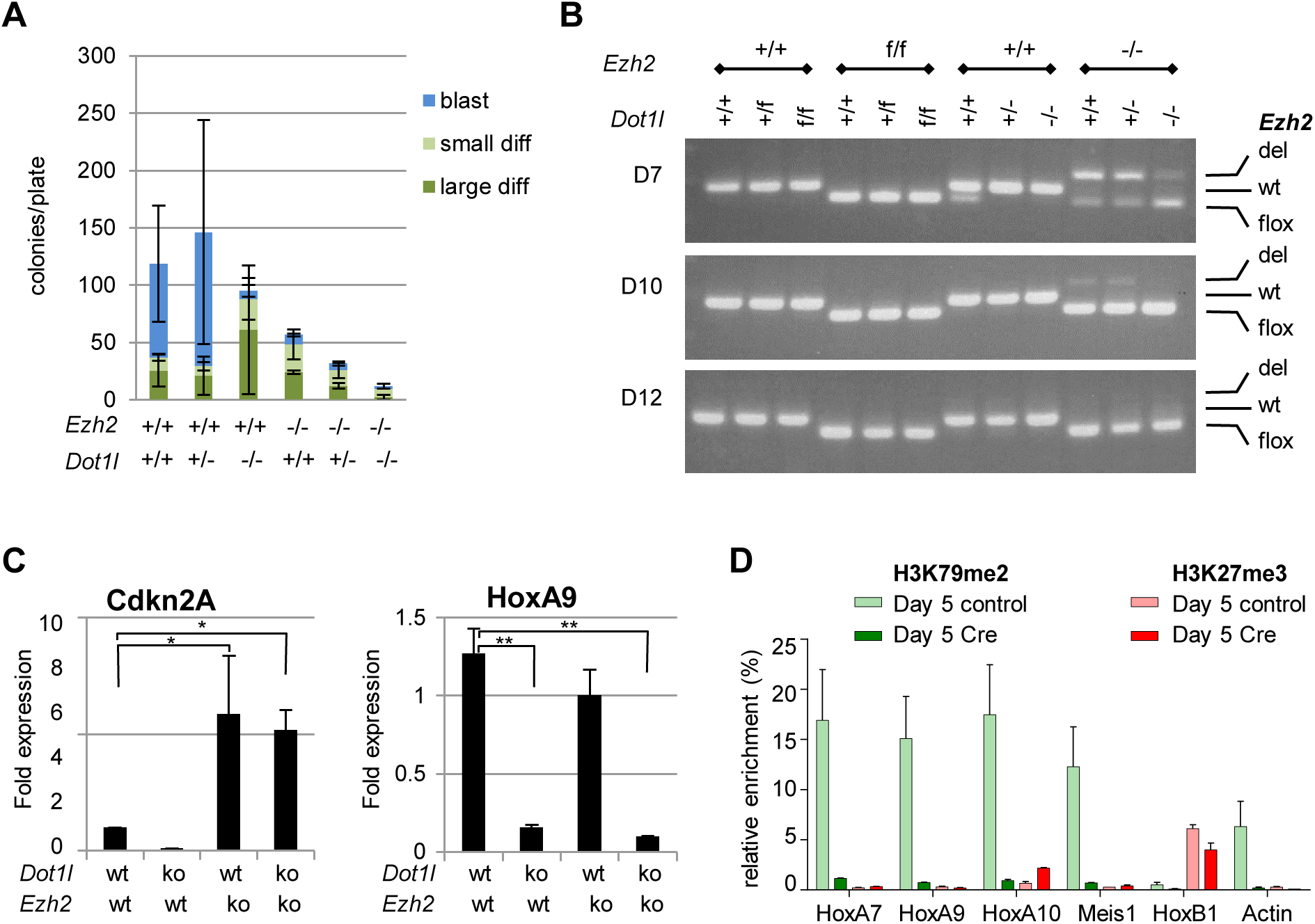
**A:** MLL-AF9 driven leukemias were established on *Dot1l* homozygous, heterozygous or wild type conditional backgrounds with either homozygous wild type or homozygous conditional *Ezh2* alleles. Deletion of conditional alleles was induced by transduction with Cre, and leukemia cells with the indicated genotypes were plated in methylcellulose. N=3 plated in duplicate, error bar: SEM, *p<0.05, **p<0.01. **B:** cells from A were maintained in liquid culture, and Ezh2 alleles were monitored at the indicated days **C:** qPCR for HoxA9 and Cdkn2A in cells from A. N=3 independent experiments, error bar: SEM, *p<0.05, **p<0.01. **D:** ChIP-qPCR over the indicated loci for H3K79me2 and H3K27me2 5 days after *Dot1l* deletion. N=2 (technical), error bars: SD.

### DOT1L and EZH2 inhibition results in both synergistic and antagonistic effects in human cell lines

Our data in the murine conditional knockout system suggest potential synergy between inactivation of DOT1L and EZH2 at high doses, mediated by de-repression of *CDKN2A*. We therefore asked whether dual inhibition of DOT1L and EZH2 would act synergistic in a panel of cell lines with intact or deleted *CDKN2A l*ocus (Figure 3A and S1, mutational information from ATCC, DSMZ and CCLE). Molm14, Monomac6, MV4;11 and THP1cells were exposed to a range of difference concentrations of DOT1L and EZH2 inhibitor alone or in combination. We observed strong synergy in Molm14 and Monomac6 cells (Figure 3B), and moderate to strong antagonism in MV4;11 and THP1 cells (Figure 3C). Thus, *CDKN2A* status did not correlate with synergy versus antagonism in the human cell lines. As in the murine system, the expression of KMT2A-fusion target genes was not affected by inhibition of EZH2. The substantial antagonistic effects between DOT1L and EZH2 inhibition in half the cell lines tested suggest that this is not a combination that should be pursued clinically.

**Figure 3:**
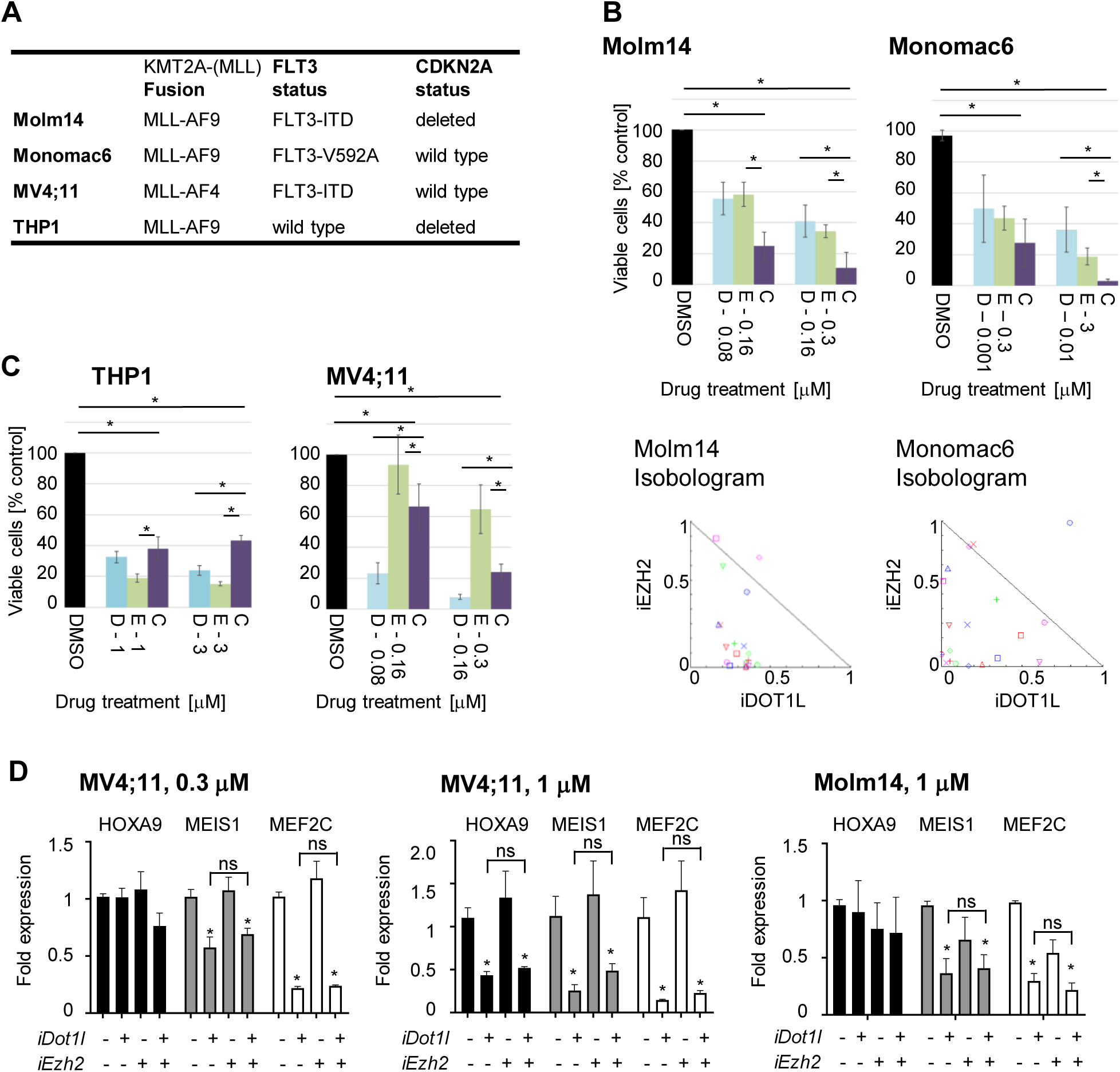
**A:** *KMT2A (MLL), FLT3* and *CDKN2A* status of Molm14, Monomac6, MV4;11 and THP1 cell lines (source: ATCC, DSMZ and CCLE). **B+C:** Cells were treated with a range of doses of EPZ5676 (“D”, DOT1L inhibitor) and GSK126 (“E”, EZH2-inhibitor) alone and in combination (“C”). Cells were replated at equal concentrations every 3-4 days. Cell numbers and viability was assess on day 14 by cell counting and trypan blue exclusion. **B** top panel: Molm14 and THP1 viable cells at the indicated dose levels. N = 3 independent experiments, error bar: SEM, *p<0.05. Bottom panel: CI-isobologram over the entire dose range (CompuSyn). **C:** THP1 and MV4;11 viable cells at the indicated dose levels. N = 3 independent experiments, error bar: SEM, *p<0.05 **D:** qPCR analysis for HOXA9 (black), MEIS1 (grey) and MEF2C (white) after exposure to of EPZ5676 (“D”, DOT1L inhibitor) and GSK126 (“E”, EZH2-inhibitor) alone and in combination at the indicated dose levels (0.3 or 1 μM) in MV4;11 and MOLM14 cells. N = 3 independent experiments, error bar: SEM, *p<0.05 compared to DMSO, ns = no significant difference between indicated samples.

### DOT1L and EZH2 inhibitors have opposing effects on the transcription of ribosomal genes

The antagonism of DOT1L and EZH2 in MV4;11 cells was particularly striking. We therefore performed RNA-Seq to investigate potential mechanisms of antagonism. Samples with dual inhibition clustered separate from single inhibition, but were closer to DOT1L than EZH2 exposed samples (Figure 4A). Principal component analysis revealed that, for the most part, DOT1L and EZH2 act on distinct, non-overlapping gene sets (Figure 4B). H3K79me2 (catalyzed by DOT1L) is associated with actively transcribed genes, while H3K27me3 (catalyzed by EZH2) is associated with silencing. In order to interrogate which pathway(s) might be responsible for the observed antagonistic effects, we therefore focused on genes that were downregulated upon DOT1L inhibition, de-repressed upon EZH2 inhibition, and normalized upon dual inhibition. The gene set in cluster 1 and 5 follows this pattern (Figure 4C+D, red and light blue cluster/line). KEGG pathway analysis showed that this gene set is highly enriched for ribosomal genes (Figure 4E). In fact, a role for PRC2 in repressing Polymerase II transcribed RNA transcription (rRNAs, tRNAs) has been previously documented [27].

**Figure 4:**
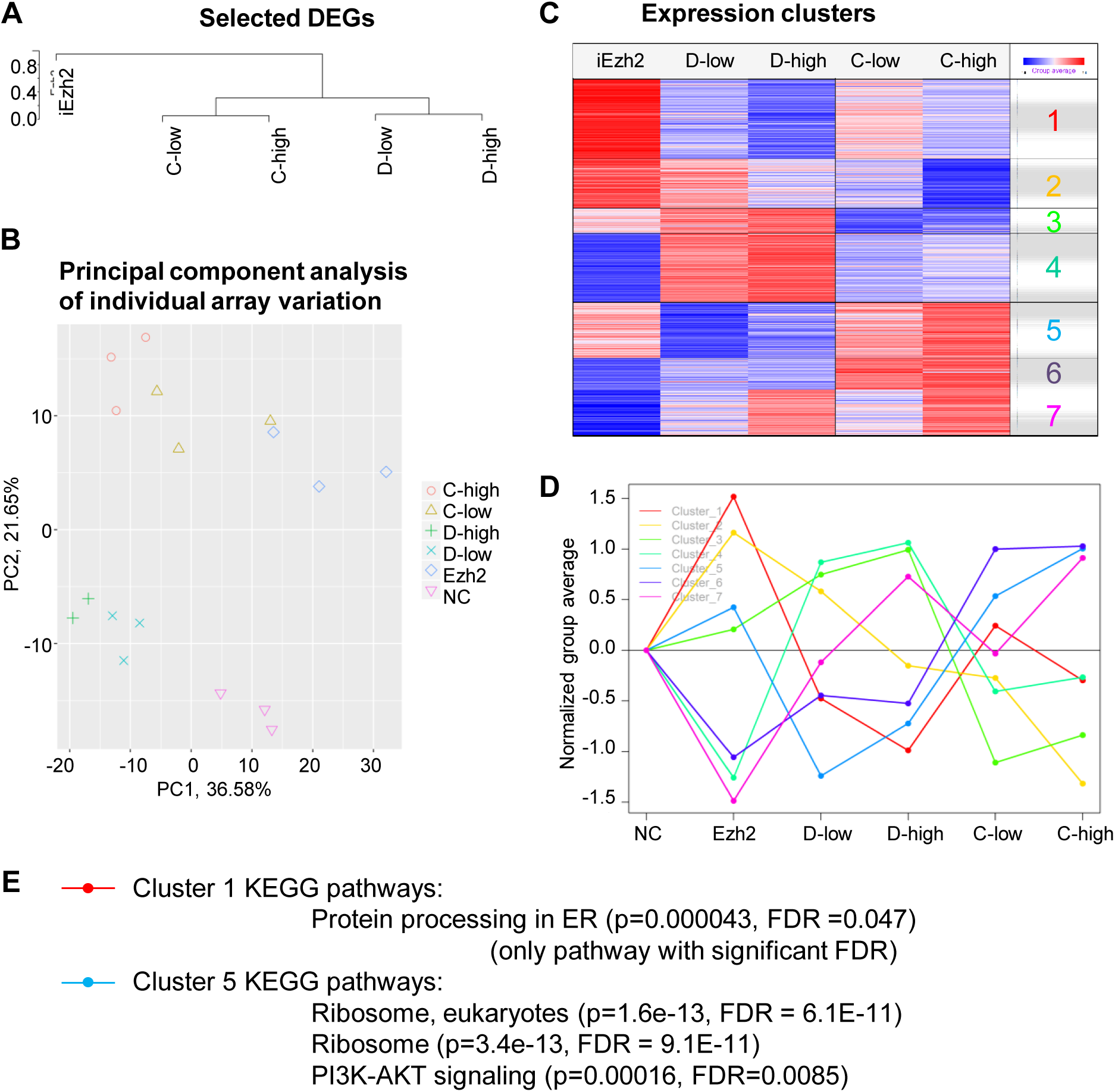
RNA-Seq of MV4;11 exposed to 0.08 μM (D-low) or 0.16 μM (D-high) DOT1L inhibitor EPZ5676 or EZH2 inhibitor GSK126 (iEzh2) alone or in combination (C-low and C-high). **A:** unsupervised clustering of drug exposed cells. **B:** Principal component analysis. **C+D:** Clusters of gene expression patterns in drug exposed cells. Cluster 1 and 5 include genes who’s expression increases upon EZH2 inhibition, decreases upon DOT1L inhibition, and is rescued by the combination. **E:** KEGG pathway analysis of cluster 1 and cluster 5 gene sets.

### DOT1L inhibition reduced protein translation

We next asked whether DOT1L inhibition reduced protein translation in MV4;11 cells. Protein translation was measured by incorporation of OP-Puro and found to be affected by DOT1L inhibition (Figure 5A). The rate of protein translation is different in different phases of the cell cycle, and we and others showed decreased cycling as one of the most prominent effects of DOT1L inhibition or deletion. We therefore chose a very early time point, and measured cell cycle distribution at the same time as protein translation. In fact, decreases in protein translation were one of the earliest effects measured when exposing MV4;11 cells to DOT1L inhibition (Figure 5A), occurring before any substantial effects on cell cycle (Figure 5B), and at a dose much lower than doses that affect HOXA cluster expression (compare to Figure 3D). EZH2 inhibition not only rescued ribosomal gene expression (Figure 4), but also partially rescued protein synthesis (Figure 5C). The antagonism between DOT1L and EZH2 inhibitors in MV4;11 cells may thus in part be driven by their opposing effects on ribosomal gene expression and, as a consequence, protein translation.

**Figure 5:**
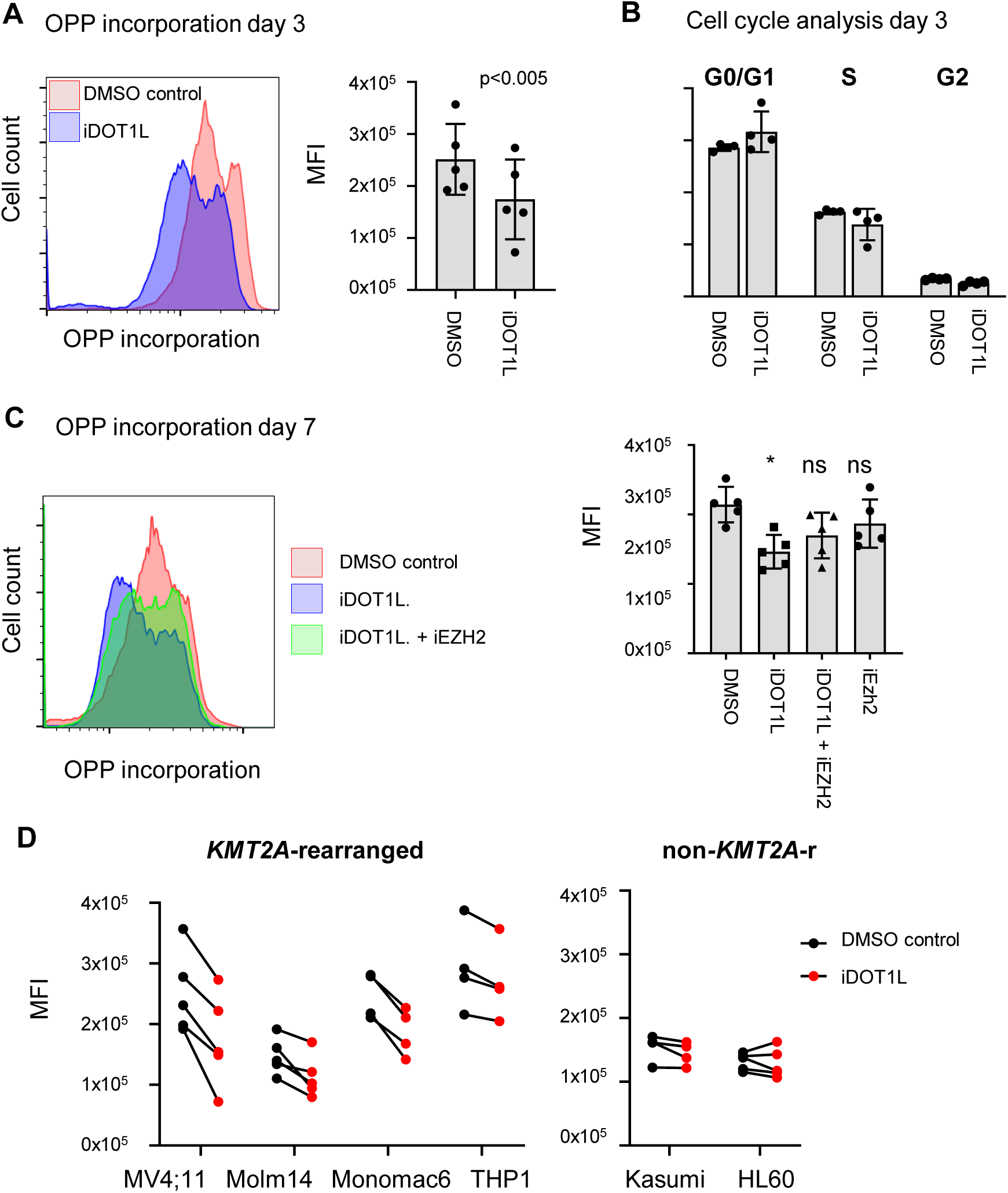
**A**: Effect of DOT1L inhibition (0.08 μM EPZ5676) on protein translation measured as OPP incorporation on day 3 after exposure in MV4;11 cells. **B:** Cell cycle analysis of MV4;11 cells on day 3 of exposure to 0.08 μM EPZ5676 **C:** Partial rescue of the effect of DOT1L inhibition on protein translation by EZH2 inhibitor. **D:** Effect of DOT1L inhibition on protein translation measured as OPP incorporation in *KMT2A*-rearranged cell lines: MV4;11 (0.08 μM), Molm14 (1 μM), THP1 (1 μM), Monomac6 (0.1 μM) (left panel, p<0.02, 2-way Anova) and non*-KMT2A*-rearranged cell lines: Kasumi (5 μM), HL60 (5 μM) (right panel, p=not significant, 2-way Anova).

**Figure 6:**
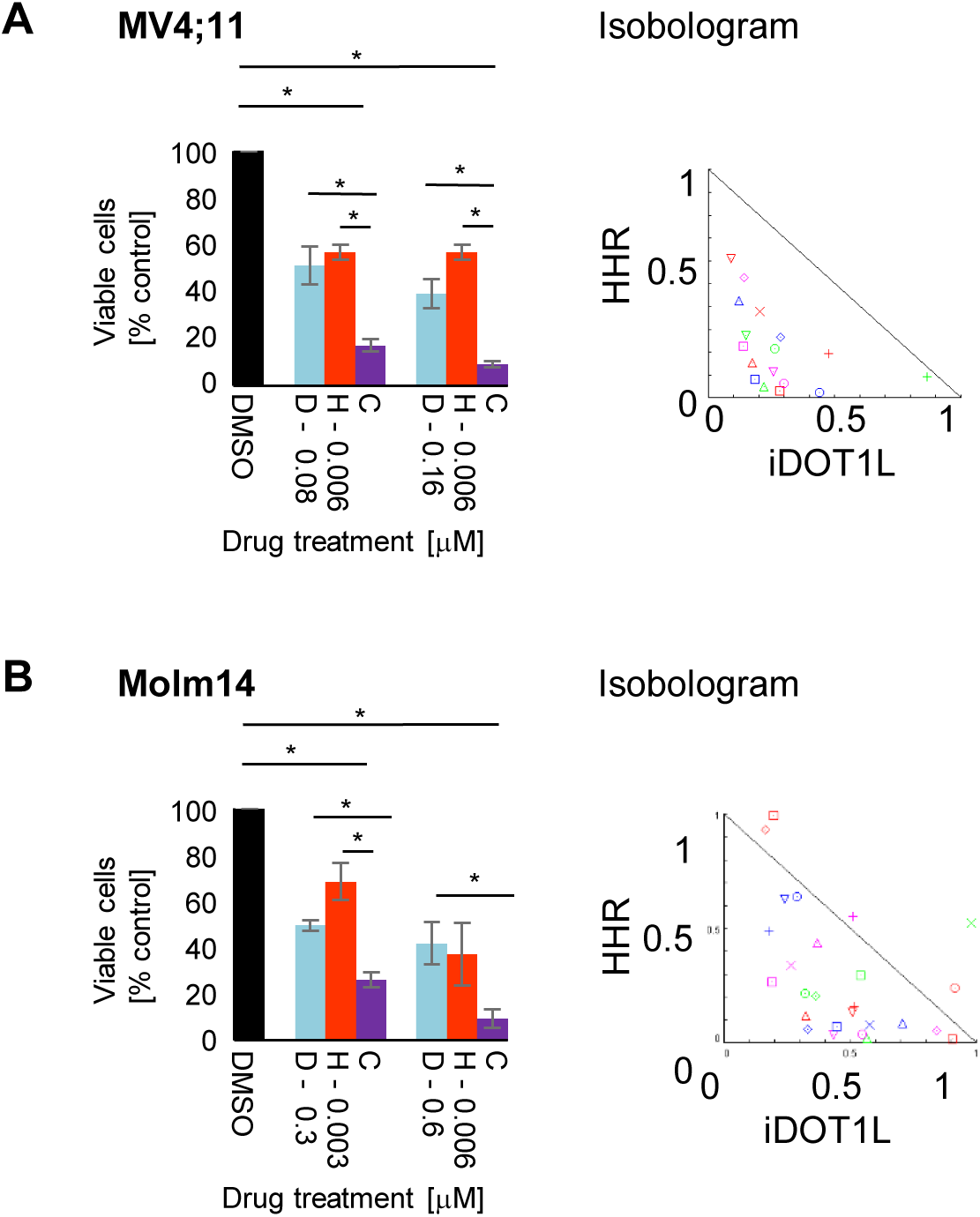
**A:** MV4;11 and. **B:** Molm14 cells were pretreated with a range of doses of EPZ5676 (“D”, DOT1L inhibitor). Cells were then exposed to Homoharringtonine (“H”) alone and in combination (“C”), and viability was assessed at 72 hours. Left panel: synergistic effects at the indicated dose levels. N=3 independent experiments, error bar: SEM, *p<0.05. Right panel: CI-isobologram over the entire dose range (CompuSyn).

### DOT1L inhibition acts synergistically with the protein translation inhibitor homoharringtonine (HHR)

The observation that DOT1L inhibition affects ribosomal gene expression and protein translation raised the possibility that EPZ5676 exposure would sensitize cells to the effect of a protein translation inhibitor. We first asked whether Inhibitory effects on protein translation were observed across all *KMT2A-*rearranged cell lines used in this study, and found this to be the case (Figure 5D),. Non-*KMT2A-*rearranged control cell lines were not affected (Figure 5D). The protein translation inhibitor homoharringtonine (omacetaxine) was FDA approved for chronic myeloid leukemia (CML) in 2012 [28, 29]. It also has published preclinical activity in several AML models and small clinical trials [30-32]. We exposed two *KM2A*-rearranged AML cell lines (MV4;11, Molm14) to 4-7 days of EPZ5676, followed by exposure to EPZ5676 and HHR. We found the dual inhibition to be synergistic, suggesting that decreased ribosomal gene expression upon DOT1L inhibition sensitized leukemia cells to the effect of a protein translation inhibitor.

## DISCUSSION

In this study, we set out to interrogate the interplay between DOT1L and PRC2 inhibition in KMT2A-rearranged leukemias. We discovered that:

### H3K27 and H3K79 methylation do not affect each other

Despite a potential rationale that modulation of H3K27 methylation might affect H3K79 methylation via binding of AF10 [19], we find no such effect over a wide range of doses of the EZH2 inhibitor GSK126. Furthermore, our results are consistent with the previously reported lack of effects of DOT1L loss or inhibition on H3K27me3 [3, 4].

### The polycomb repressive complex 2 is not responsible for silencing the HOXA cluster in KMT2A-fusion driven leukemia

Despite a clear role of PRC2 mediated silencing of the HOXA cluster in other contexts, the HOXA cluster does not acquire H3K27 methylation in a time frame consistent with a primary silencing mechanism. Furthermore, dual inhibition/inactivation of DOT1L and EZH2 does not rescue HOXA cluster expression. These results were consistent across the murine retroviral model as well as human cell lines. Given the well documented regulation of the HOXA by PRC2 during development and in the context of other subtypes of leukemia, our data is highly relevant in that is suggests a more complex and context dependent regulation of the HOXA cluster than previously appreciated.

### The polycomb repressive complex 2 is not responsible for silencing the CDKN2A in human *KMT2A-*rearranged cell lines

CDKN2A is a major canonical target of PRC2. Although *CDKN2A* independent effects of loss of PRC2 on leukemogenesis have been reported [33], de-repression of *CDKN2A* was described as a major mechanism for the anti-leukemic effect of PRC2 inactivation [7, 9]. In the murine model, persistent downregulation of the KMT2A-fusion target HOXA9 and simultaneous de-repression of *CDKN2A* provide a good potential mechanism for the observed synergy between inactivation of DOT1L and EZH2. However, synergistic versus antagonistic effects in a panel of *human* cell lines did not correlate with *CDKN2A* status, and in cells with an intact *CDKN2A* locus no de-repression was observed upon inhibition of EZH2.

### Inhibition of DOT1L affects ribosomal gene expression, protein translation, and sensitized *KMT2A*-rearranged cell lines to homoharringtonine

More in depth analysis of the antagonistic effects between DOT1L and EZH2 inhibition in MV4;11 cells revealed that EZH2 inhibition mediates resistance to DOT1L inhibition through partial rescue of ribosomal gene expression and protein translation. With respect to EZH2, our data is consistent with the previously reported effect of PRC2 on Polymerase III transcribed non-translated RNA gene transcription [27]. Furthermore, we found that DOT1L inhibition resulted in decreased protein synthesis in *KMT2A*-rearranged AML cell lines. This resulted in sensitization to the effect of the protein translation inhibitor HHR. A wealth of preclinical data as well as several clinical trials [30-32] support HHR as an active agent in AML. Our data suggests that the combination of DOT1L inhibition with HHR merits further investigation, with the potential for rapid clinical translation given that HHR is FDA.

## ACKNOWLEDGEMENTS

We thank Vikram Paralkar at the University of Pennsylvania for helpful critical discussions. This work was supported by start-up funds from University of Colorado Denver, Hematology/Oncology Section and the Children’s Hospital Colorado Research Institute (KMB and TN), start-up funds by the Division of Pediatric Oncology and the Abramson Cancer Research Center at the Children’s Hospital of Philadelphia (KMB), as well as NHLBI K08HL102264 (KMB).

## AUTHOR CONTRIBUTIONS

KMB and TN designed the study, AL, SSR, NZ and KMB performed the experiments and analyzed data, ZFY performed and analyzed mass spectrometry, HMX performed bioinformatic analysis, AL and KMB wrote the manuscript with input from all co-authors.

