## Supplemental Figure for "Epigenetic regulation of protein translation in *KMT2A*-rearranged AML"

Supplemental Figure 1

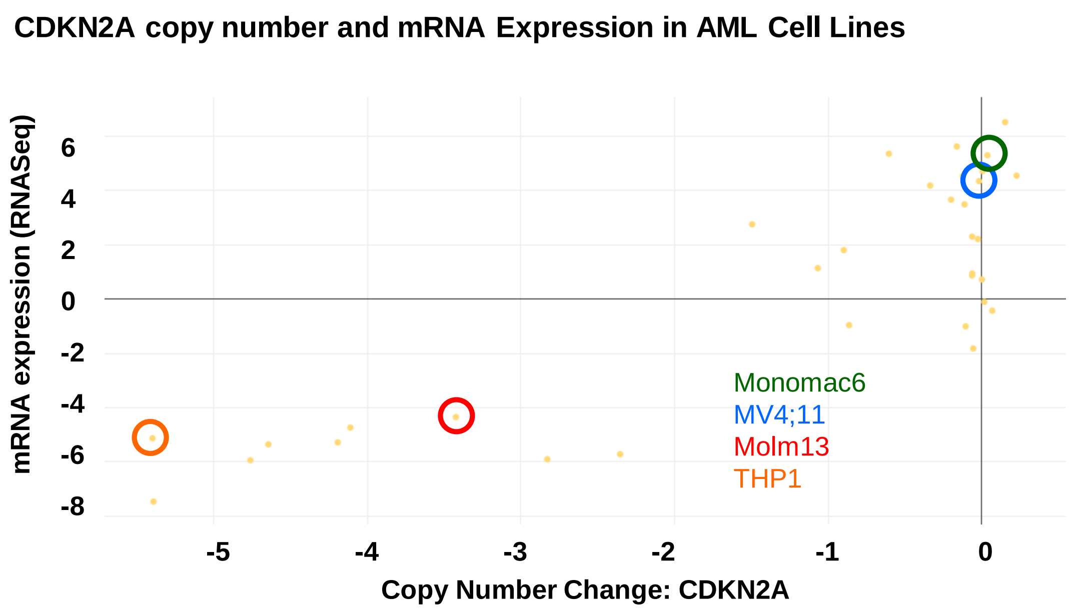


**Supplemental Figure 1**
CDKN2A copy number and expression in leukemia cell lines, the cell lines used in this study (Monomac 6, MV4;11, Molm13, THP1) are highlighted. Source: Broad Insititue Cancer Cell Line Encyclopedia (CCL): https://portals.broadinstitute.org/ccle
