## Supplemental Materials and Methods for "Epigenetic regulation of protein translation in *KMT2A*-rearranged AML"

| Reagent or Resource | Source | Identifier |
|---|---|---|
| **Western Blots** | | |
| Rabbit polyclonal anti-H3K79me2 | Abcam | Cat # ab3594 |
| Anti-Histone H3 antibody | Abcam | ab1791 |
| Goat anti rabbit secondary H+L-HPR conjugated | BioRad | Cat# 170-6515 |
| 10% BisTris Gel | Invitrogen | Cat# NP0301 |
| Western Lighting RTM Plus-ECL | Perkin-Elmer | Cat# NEL104001EA |
| **Flow Cytometry and Sort** | | |
| CD8α, Biotin, anti-mouse | Biolegand | Clone 53-6.7<br>Cat# 100704 |
| Gr1 (Ly6-G/Ly6-C), Biotin, anti-mouse | Biolegand | Clone RB6-8C5<br>Cat# 108404 |
| B220, Biotin, anti-mouse/human | Biolegand | Clone RA3-6B2<br>Cat# 103204 |
| CD19, Biotin, anti-mouse | Biolegand | Clone 6D5<br>Cat# 115504 |
| IL-7Rα (CD127), Biotin, anti-mouse | Biolegand | Clone A7R34<br>Cat# 135006 |
| Ter-119, Biotin, anti-mouse | Biolegand | Clone TER-119<br>Cat# 116204 |
| Streptavadin, APC-Cy7 | Biolegand | Cat# 405208 |
| Ckit (CD117), Alexa Fluro 647, anti-mouse | Biolegand | Clone 2B8<br>Cat# 105818 |
| Sca-1 (Ly6A), Pe-Cy7, anti-mouse | Invitrogen | Clone D7<br>Ref# 25-5981-82 |
| CD2, APC, anti-human | Biolegand | Clone RPA-2.10<br>Cat# 300214 |
| Dyanbeads M-280, Streptavidin | Invitrogen | Ref# 11206D |
| APC annexinV | BD Biosciences | Cat# 550474 |
| DAPI | BD Biosciences | Cat# 564907 |
| **Plasmids** | | |
| MSCV-MLL-AF9-GFP | Armstrong Lab | NA |
| Cre-IRES-pTomato (Cre) | Armstrong Lab | NA |
| MSCV-IRES-pTomato | Armstrong Lab | NA |
| **Cytokines** | | |
| Recombinant Murine IL-3 | PeproTech | Cat# 213-13 |
| Recombinant Murine IL-6 | PeproTech | Cat# 216-16 |
| Recombinant Murine SCF | PeproTech | Cat# 250-03 |
| Recombinant Murine TPO | PeproTech | Cat# 315-14 |
| Recombinant Murine FLT3-Ligand | PeproTech | Cat# 250-31L |

| | | |
|---|---|---|
| **Cell Culture Reagents** | | |
| RPMI-1640 Media | VWR | Cat# 10-040-CV |
| IMDM (medium iscoves modif of DMEM) | VWR | Cat# 45000-366 |
| DMEM | VWR | Cat# 45000-312 |
| M3234 | | |
| Fetal Bovine Serum | Life Technologies | Cat# 10438026 |
| L-Glutamine | Life Technologies | Cat# 25030081 |
| Penicillin-Streptomycin (10,000 U/mL) | Invitrogen | Cat# 15140122 |
| Fugene 6 Transfection Reagent | VWR | Cat# PAE2692 |
| Optimem | Thermo | Cat# 31985-062 |
| Retronectin | Clontech Laboratories | Cat# T100B |
| Poly(ethylene glycol) | Sigma | Cat# P4338 |
| Trypan Blue Solution | Mediatech | Cat# MT25-900-CI |
| Phosphate Buffered Saline | Mediatech | Cat# MT21-031-CV |
| BD Pharmlyse | Fisher BD | Cat# 555899 |
| **Chemicals** | | |
| TritonX-100 | VWR | Cat# 9002-93-1 |
| Hydrochloric Acid | Fisher Scientific | Cat# A144 |
| Sodium Chloride (NaCl) | Sigma Aldrich | Cat# S9888 |
| NP-40 (IGEPAL) | Alfas Aesar | Cat# J61055 |
| Sodium dodecyl sulfate (SDS) | Teknova | Cat# S0288 |
| Sodium Deoxycholate | Alfas Aesar | Cat# J62288 |
| Tris-HCl | Roche | Cat # 10812846001 |
| Lithium Chloride (LiCl) | Alfas Aesar | Cat# 36217 |
| Ethylenediaminetetraacetic acid (EDTA) | VWR | Cat# E1777 |
| Sodium Bicarbonate ($NaHCO_3$) | Fisher Scientific | Cat# S233 |
| XTT assay | Sigma | Cat#11465015001 |
| **Kits** | | |
| DNA and RNA Extraction Kits | | |
| QiAquick PCR Purification Kit | Qiagen | Cat# 28106 |
| RNeasy Plus Mini Kit | Qiagen | Cat# 74136 |
| RNeasy Plus Micro Kit | Qiagen | Cat# 74034 |
| ZymoPURE II Plasmid Maxi Prep Kit | Zymo | Cat# 11-555B |
| Click-iT EdU Alexa Fluor™ 647 Flow Cytometry Assay Kit | Invitrogen | Cat# C10419 |
| Click-iT™ Plus OPP Alexa Fluor™ 488 Protein Synthesis Assay Kit | Invitrogen | Cat# C10456 |

| **Genotyping and Deletion PCR Primers** | |
|---|---|
| Primer Name | 5'→3' Sequence |
| DOT1L 6F | GGTGCGACTTCTGGAACCTA |
| DOT1L 6R | CACCGAAAAACAATGTCTCG |
| DOT1L 1R | TCTTTGAGCCGACCTTGAGT |
| EZH2 F1 | ACTTACTGCTGGCACCGTCT |
| EZH2R1 | AAGTCAACTGGAATGCACGA |
| **qPCR Primers /tests** | |
| huHOXA9_qPCR_F | CTGTCCCACGCTTGACACTC |

| | |
|---|---|
| huHOXA9_qPCR_R | CTCCGCCGCTCTCATTCTC |
| huMeis1_F | GGGCATGGATGGAGTAGGC |
| huMeis1_R | GGGTACTGATGCGAGTGCAG |
| huMef2c_qPCR_F_1 | GAACGTAACAGACAGGTGACAT |
| huMef2c_qPCR_R_1 | CGGCTCGTTGTACTCCGTG |
| huGAPDH | GAAGGTGAAGGTCGGAGTC |
| huGAPDH | GAAGATGGTGATGGGATTTC |
| mHOXA9 | 4331182-Mm00439364_m1 (Applied Biosystems) |
| mCDKN2A_F | 4331182-Mm00494449_m1 (Applied Biosystems) |
| mGAPDH | 4331182-Mm99999915_g1 (Applied Biosystems) |
| mACTB | 4331182-Mm01205647_g1(Applied Biosystems) |
| mHOXA7_ChIP_F | CTCTTCTGTTTCCCATCCTGGT |
| mHOXA7_ChIP_R | GGCAAT ATCCGGGATCCACT |
| mHOXA9_ChIP_F | GAATAGGAGGAAAAAACAGAAGAGG |
| mHOXA9_ChIP_R | TGTATGAACCGCTCTGGTATCCTT |
| mHOXA10_ChIP_F | CCTTTTTGGTCGACTCGCTC |
| mHOXA10_ChIP_R | CAACACCAGCCTCGCCTCT |
| mMeis1_ChIP_F | TCACCACGTTGACAACCTCG |
| mMeis1_ChIP_R | GCTTTCTGCCACTCCAGCTG |
| mHOXB1_ChIP_F | GGGACTGCCAAACTCTGGC |
| mHOXB1_ChIP_R | CATGTGATCTCTCCCAGGCC |
| mActin_ChIP_F | GGGAACCAGACGCTACGATC |
| mActin_ChIP_R | TTGGACAAAGACCCAGAGGC |

**Supplemental Methods:**

**Cell lines:** Human leukemia cell lines (MV4;11, Molm14, Monomac6, THP1, Kasumi, HL60, 293T) were obtained from Deutsche Sammlung von Mikroorganismen und Zellkulturen (DSMZ, Germany) or the American Type Culture Collection. Cells were maintained in RPMI-1640 media supplemented with heat-inactivated 10% fetal bovine serum, 50 U/ml Penicillin/Streptomycin. 293T cells were maintained in DMEM supplemented with heat-inactivated 10% fetal bovine serum, 50 U/ml Penicillin/Streptomycin. All cells were cultured in the appropriate media a humidified incubator at 37°C in 5% CO2. Cells were tested for mycoplasma and re-authenticated every 6 months in culture.

**Drug Assays**: EPZ4777 was obtained from Epizyme. EPZ5676, GSK126 and HHR were obtained from Cayman chemical. Compounds were dissolved in DMSO, and all dilution series were prepared keeping the DMSO exposure euqal accross all conditions and at <0.1% (<1:1000). Human leukemia cell lines were exposed to EPZ5676 (DOT1L inhibitor), GSK 126 (EZH2 Inhibitor) or homoharringtonine at the indicated concentrations, and cells were replated at equal densities in fresh compound containing media every 3-4 days. Cell growth and viability was assessed either by serial replating and trypan blue exclusinon, or XTT assay.

***Dot1l* and *Ezh2* knockout mice, breeding:** Animals were maintained at the Animal Research Facility at the Boston Children's Hospital. Animal experiments were approved by the Internal Animal Care and Use Committee*. Dot1l* [3] and *Ezh2 ([7])* conditional knockout mice were previously described and were maintained on a C57BL/6 background.

**Generation of transformed murine cells:** Ecotropic retroviral vectors containing murine KMT2A-MLLT3-IRES-GFP, Cre-IRES-pTomato (Cre) and MSCV-IRES-pTomato (MIT) were generated by cotransfection of 293 cells. Bone marrow cell suspensions from mice conditional for *Dot1l* and/or *Ezh2* and litter mate controls were prepared by isolating whole bone marrow and performing red cell lysis with Pharm Lyse. Lineage depletion was performed using biotinylated monoclonal antibodies to CD3e, CD4, CD8a, CD19, B220, Gr-1, IL-7R and Ter-119. Cells were subjected to 2 rounds of magnetic bead depletion with streptavidin conjugated Dynabeads. Lineage depleted (lin- ) cells were stained with with APC-Cy7 conjugated streptavidin c-Kit Alexa 647, Sca-1 PE-Cy7 and sorted for Lin- Sca-1+ cKit+ (LSK). Sorted cells were pre-stimulated for 24 h with 10 ng/ml mIL3 and mIL6 and 20 ng/ml mSCF, mFlt3L and TPO. Transduction was carried out on retronectin with MLL-AF9 in the presence of murine IL3, IL6, SCF, Flt3L and TPO in concentrations as above.

**Biochemical Assays (cell growth, colony growth, apoptosis, cell cycle, western blotting):** For colony assays, sorted transduced cells were plated in methylcellulose M3234 containing IL3, IL6 and SCF at 1000 cells per plate in duplicate, and colonies were scored after 7 days of culture. For liquid culture of murine cells, cells were maintained in media with cytokine support, and replated every 2-3 days. *Dot1l* and *Ezh2* deletion was verified by PCR at each replating beyond day 7. Cell growth and viability were followed by serial cell counts using Trypan blue exlcusion. Apoptosis and cell cycle analysis were performed using the Annexin-staining and the Click-IT EdU kit. Protein translation was analyzed using the Click-IT OP-Puro kit. Western blotting for histone modifications was performed on purified histones isolated by acid extraction using the indicated antibodies and controls.

**Histone Mass Spectrometry:** Histones were isolated, chemically derivatized and analyzed by mass spectrometry as previously described [19].

**qPCR analysis of HOXA9 and CDKN2A:** RNA was isolated from sorted murine transformed progenitor cells or compound treated human cell lines using RNeasy mini columns (Qiagen). Please refer to the table above for primer sequences / tests. Fold-change of is shown compared vehicle treated control (human cell lines) or wild type control (murine knockout studies).

**Chromatin immunoprecipitation (ChIP):** Chromatin immunoprecipitation for H3K79me2 and H3K27me3 in murine KMT2A-MLLT3 leukemias was performed using rabbit polyclonal antibodies from abcam (ab3594 Cambridge, MA) similarly as described [20]. Briefly, cells were fixed in PBS 1% formalin (v/v) with gentle rotation for 10 minutes at room temperature. Fixation was stopped by the addition of glycine (125 mM final concentration). Fixed cells were washed twice in ice-cold PBS, resuspended in SDS lysis buffer (1% SDS, 10mM EDTA, 50 mM Tris-HCl, pH 8.1). Chromatin was sheared by sonication to about 100-300bp fragments using EpiShear (Active Motive, Carlsbad, CA) and diluted tenfold with dilution buffer (0.01% SDS, 1.1% Triton-X100, 1.2 mM EDTA, 16.7 mM Tris-HCl, pH 8.1,167 mM NaCl). Magnetic bead labeled antibodies against specific histone modifications were used to precipitate DNA fragments associated with modified histones. Precipitates were washed ChIP DNA was analyzed by qPCR.

**RNA amplification and RNA-Seq:** RNA was isolated from MV4;11 cells exposed to 5 days of EPZ5676 or GSK126 using RNeasy mini columns (Qiagen). RNA for RNA-Seq was submitted to the UC-Denver genomics core for library preparation and sequencing.

**Data analysis and statistical methods**:

**Histone PTMs** were analyzed using the EpiProfile 2.0 computational algorithm [21].

**Drug interactions** were evaluated for synergy or antagonism using CompuSyn (http://www.combosyn.com/) [22, 23].

**RNA-Seq** raw Fastq files were aligned using STAR [24] against Human GRCh37 reference using default parameters and quantified by applying Kallisto [version 0.45.0, PMID: 27043002]. Output from Kallisto was then directly imported into DESeq2 [25] in order to detect differentially expressed genes (DEG). DEGs were deemed as genes with False Discovery Rate (FDR) less than 0.05 level. For the clustering analysis, we first created a hierarchical tree based on gene-gene expression level correlations crossing all treatment conditions. We cut the tree at height level 1.5. This would group genes into clusters. Obtained initial clusters were refined by calculating correlation of each gene's expression patterns to the centroid (median) of its cluster. We removed genes if the correlation is less than 0.5. We also remove clusters with size less than 10% of the expected size (estimated by number of total genes divided by number of clusters). If less than six clusters were left, reduce the height cutoff by 0.05 and repeated this step until we got enough clusters. We merged clusters if the correlation of their centroids is greater than or equal to 0.5. We iterate the step until there is no two clusters were correlated at 0.5 or more level. We further refined obtained clusters by removing genes from a cluster if its correlation coefficient to the cluster is less than 0.5 or the correlation coefficient to any other cluster is at least 0.2. We then update the clusters centroid profile. We repeated re-clustering steps for 50 times or until the re-clustering converged. Finally, we removed clusters with number of genes less than 1% of total number of unique DEGs. All analysis was carried using R, version 3.5.0
